# Application of high dimensional flow cytometry and unsupervised analysis to define the immune cell landscape of early childhood respiratory and blood compartments

**DOI:** 10.1101/2021.03.21.436363

**Authors:** Shivanthan Shanthikumar, Sarath C. Ranganathan, Richard Saffery, Melanie R. Neeland

**Author notes:** **Corresponding author:** Dr Melanie Neeland, Murdoch Children’s Research Institute, Royal Children’s Hospital, Parkville, Victoria, Australia, 3052, P:^+^61419573622, E. **Author Contributions:** S.S, S.C.R, R.S and M.R.N contributed to the concept and design of the study. S.S. and S.C.R were responsible for patient recruitment and sample collection. S.S. and M.R.N performed experiments, analysed the data and co-wrote the manuscript. M.R.N supervised the work and interpreted the findings. M.R.N, R.S, and S.C.R. provided funding. All authors edited and approved the final manuscript.

## Abstract

The cellular landscape of the paediatric respiratory system remains largely uncharacterised and as a result, the mechanisms of highly prevalent childhood respiratory diseases remain poorly understood. A major limitation in defining mechanisms of disease has been the availability of tissue samples collected in early life, as well as technologies that permit deep immune analysis from limited sample volumes. In this work, we developed new experimental methods and applied unsupervised analytical tools to profile the local (bronchoalveolar lavage) and systemic (whole blood) immune response in childhood respiratory disease. We quantified and comprehensively phenotyped immune cell populations across blood and lung compartments in young children (under 6 years of age), showed that inflammatory cells in the BAL express higher levels of activation and migration markers relative to their systemic counterparts, and applied new analytical tools to reveal novel tissue-resident macrophage and infiltrating monocyte populations in the paediatric lung. To our knowledge, this is the first description of the use of these methods for paediatric respiratory samples. Combined with matched analysis of the systemic immune cell profile, the application of these pipelines will increase our understanding of childhood lung disease with potential to identify clinically relevant disease biomarkers.

## INTRODUCTION

A detailed understanding of the tissue-specific immune landscape in health and disease is required to improve the clinical management of many childhood diseases. Aberrant inflammation is a hallmark of several childhood lung diseases, including bronchopulmonary dysplasia (Balany and Bhandari, 2015), preschool wheeze (Xepapadaki et al., 2020), asthma (Chedevergne et al., 2000), cystic fibrosis (Tirouvanziam et al., 2000), primary ciliary dyskinesia (Cockx et al., 2017), and COVID-19 (Neeland et al., 2021). Despite this, little is known regarding the immune cell profiles and mechanisms governing inflammatory processes in the early life respiratory system.

A major limitation in defining immune cell development in the paediatric lung has been the availability of tissue samples collected in early life, as well as technologies that permit deep immune analysis from limited sample volumes. Unlike adults, children infrequently undergo surgical procedures for evaluation of lung diseases, and as such research samples are not readily obtained. One clinical test which can be leveraged for research purposes in children is the bronchoalveolar lavage (BAL), which samples immune cells in the lung.

Furthermore, the recent advancement of multiple single cell technologies, including single-cell RNA sequencing (sc-RNAseq), single cell DNA methylation analysis (Karemaker and Vermeulen, 2018), and single cell Assay for Transposase-Accessible Chromatin sequencing (Buenrostro et al., 2015), along with improvements in existing techniques such as flow cytometry and CyTOF, now mean that small volumes of childhood BAL fluid (collected at the time of clinically indicated procedures) can be used to profile the immune cells of the lung in highly granular detail.

These technologies are complemented by new unsupervised analysis tools, including hierarchical clustering (e.g. Seurat (Hao et al., 2020), FlowSOM (Van Gassen et al., 2015)) and dimensionality reduction (e.g. UMAP (Konopka, 2018), tSNE (Laurens van der Maaten, 2008)) algorithms that reduce reliance on prior knowledge and assumptions, and permit unbiased assessment of high dimensional data. Consequently, unsupervised analyses are increasingly viewed as the gold standard assessment of single cell data

We previously published a flow cytometry-based protocol for immune phenotyping of early life BAL fluid (Shanthikumar et al., 2020). This work revealed that the most common immune cell population in the paediatric lung is the alveolar macrophage, comprising up to 90% of immune cells in BAL. This was followed by granulocytes, making up approximately 5% of immune cells, as well as small proportions of lymphocytes, monocytes, and dendritic cells. Whilst the first to report detailed immune cell frequencies from childhood lung samples, there were several limitations of this work, including use of cryopreserved samples (resulting in incomplete phenotyping of granulocytes), as well as limited unsupervised analyses.

In the present study, we sought to provide advanced experimental and analytical methods for immune cell profiling of early life respiratory disease. We performed simultaneous assessment of BAL fluid and whole blood for comparisons between circulating and tissue-resident immune cell profiles, analysed functional markers associated with cell activation and migration, and applied new unsupervised clustering and visualisation analysis tools to both datasets. This is the first description of the use of these methods for paediatric respiratory samples and highlight their utility in developing a better understanding of the role of inflammation in lung diseases of childhood.

## MATERIALS AND METHODS

### Study participants

All subjects have cystic fibrosis (CF) and are enrolled in the AREST CF cohort (Sly et al., 2013). Table 1 describes the demographics of the study participants. All CF subjects were cared for at the Royal Children’s Hospital, Melbourne, Australia. Their families gave written and informed consent for their involvement in the AREST CF research program (HREC #25054), which includes collection of samples and clinical data.

**Table 1.** Demographics of the study cohort

| Patient | Sex | Age<br>(months) | BAL viability<br>(% single cells) | Blood viability<br>(% single cells) |
| --- | --- | --- | --- | --- |
| 1 | F | 27 | 88.8 | 98.9 |
| 2 | M | 60 | 76.7 | 98.3 |
| 3 | F | 54 | 78.7 | 99.8 |
| 4 | F | 55 | 82.6 | 98 |
| 5 | F | 24 | 81 | 99.8 |
| 6 | M | 62 | 80.4 | 99.7 |
| 7 | M | 54 | 82.9 | 97.8 |
| 8 | M | 23 | 73 | 96.2 |
| 9 | M | 24 | 74 | 99.8 |
| 10 | M | 17 | 77.7 | 98.8 |
| 11 | M | 11 | 72 | 97.4 |

### Sample collection

As part of routine care, subjects underwent annual BAL around the time of their birthday from 1-6 years inclusive. BAL was performed under general anaesthesia. Each BAL aliquot consisted of 1mL/kg (maximum 20mL) of normal saline being inserted via the working channel of the bronchoscope and then suctioned for return. BAL samples were placed on ice and processed for flow cytometry within 1 hour of the procedure. Venous blood samples were collected in EDTA tubes from each participant at the time of BAL collection. Blood samples were kept at room temperature and processed for flow cytometry within 1 hour of collection.

### Flow cytometry of BAL and whole blood

BAL samples were centrifuged at 300 x g for 10 mins at 4°C. Cell-free BAL supernatant was then collected and stored at −80°C. The cell pellet was resuspended in 10mL of media (RPMI supplemented with 2% fetal calf serum (FCS)), filtered through a 70uM filter and centrifuged at 300 x g for 10 mins at 4°C. Supernatant was discarded and the cell pellet resuspended in PBS for viability staining using near infra-red viability dye according to manufacturers’ instructions. Following blood collection, 100 μl of EDTA whole blood was aliquoted for flow cytometry analysis and lysed with 1mL of red cell lysis buffer for 10 mins at room temperature. Cells were washed with 1mL PBS and centrifuged at 400 x g for 5 mins. Following another wash, cells were resuspended in PBS for viability staining using near infra-red viability dye according to manufacturers’ instructions. For both BAL and whole blood samples, the viability dye reaction was stopped by the addition of FACS buffer (2% heat-inactivated FCS in 2mM EDTA) and cells were centrifuged at 400 x g for 5 mins. Cells were then resuspended in human FC-block according to manufacturers’ instructions for 5 minutes at room temperature. The antibody cocktail (Supplementary Table 1) made up at 2X concentration was added 1:1 with the cells and incubated for 30 minutes on ice. Following staining, cells were washed with 2 mL FACS buffer and centrifuged at 400 x g for 5 minutes. Cells were then resuspended in 2% PFA for a 20 minute fixation on ice, washed, and resuspended in 150μl FACS buffer for acquisition using the BD LSR X-20 Fortessa and BD FACS DIVA V 9.0 software. For all flow cytometry experiments, compensation was performed at the time of sample acquisition using compensation beads. Supplementary Figure 1 depicts the manual gating strategy for BAL samples. Supplementary Figure 2 depicts the manual gating strategy for whole blood samples. Average viability as determined by viability dye for BAL and blood samples were 78.9% and 98.5%, respectively (individual sample viability is provided in Table 1).

### Data analysis

Results were analysed (manual gating, FlowSOM, UMAP) using FlowJo Version 10.7.1 software. FlowSOM and UMAP analyses of BAL and blood data were conducted using a concatenated file for each sample type containing 9,500 randomly selected live single CD45^+^ cells per individual. Algorithms were run using default parameters within FlowJo. Manually gated results are presented as proportion of CD45^+^ leukocytes. Data were plotted in Prism version 8.0.0. Boxplots show the medians, the 1^st^ and 3^rd^ quartile as well as the smallest and largest values as whiskers. Individual data points are shown. The heatmap depicts normalised cell proportions in matched BAL and blood samples for each individual. For correlations, two-sided spearman’s rank tests were performed.

## RESULTS

### Immune cell proportions in paediatric BAL and whole blood by manual gating

Matched BAL and blood samples from 11 children with CF aged between 1-5 years were used in this study (Table 1). A single flow cytometry panel consisting of 16 antibodies was used for both BAL and whole blood (Supplementary Table 1). Manual gating analysis using the gating strategies outlined in Supplementary Figures 1 and 2 was used to determine the cellular immune profile of paediatric respiratory and blood compartments (Figure 1A and B).

**Figure 1.**
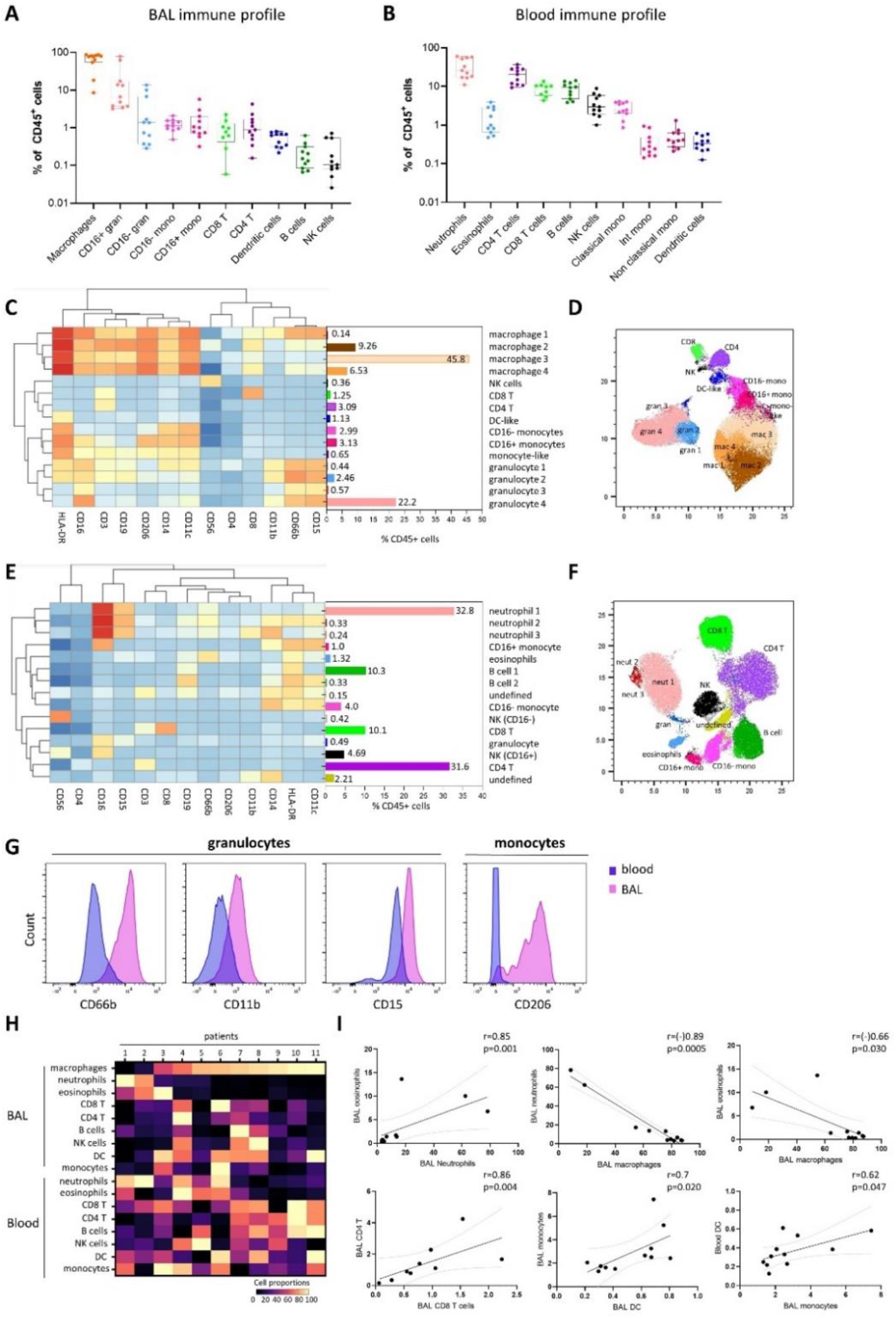
Immune cell landscape of paediatric respiratory and blood compartments. **(A)** Frequencies of macrophages, granulocytes, monocytes, CD8 T cells, CD4 T cells, dendritic cells, B cells and NK cells in BAL, expressed as proportion of CD45+ immune cells. **(B)** Frequencies of neutrophils, eosinophils, CD4 T cells, CD8 T cells, B cells, NK cells, monocytes and dendritic cells in whole blood, expressed as proportion of CD45+ immune cells. **(C)** FlowSOM clustering of BAL using lineage markers HLA-DR, CD16, CD3, CD19, CD206, CD14, CD11c, CD56, CD4, CD8, CD11b, CD66b, CD15 revealed 15 cell clusters which have been annotated as cell populations based on expression pattern and projected onto a **(D)** UMAP plot with colours corresponding to the annotated FlowSOM clusters. **(E)** FlowSOM clustering of whole blood using lineage markers CD56, CD4, CD16, CD15, CD3, CD4, CD19, CD66b, CD206, CD11b, CD14, HLA-DR, CD11c revealed 15 cell clusters which have been annotated as cell populations based on expression pattern and projected onto a **(F)** UMAP plot with colours corresponding to the annotated FlowSOM clusters. **(G)** Expression profiles of granulocytes and CD16+ monocytes in BAL and blood. **(H)** Heatmap depicting immune cell proportions in matched BAL and blood samples from all individuals in the cohort. **(I)** Line plots depicting significant correlations between immune cell populations in BAL and blood for all individuals in the cohort. Two-sided spearman’s rank correlations were performed.

For BAL, alveolar macrophages were the predominant immune cell type at a median of 76.9% of CD45^+^ leukocytes. The BAL granulocyte fraction was subtyped into CD16^+^ and CD16^−^ populations, with the CD16^+^ granulocytes comprising a median of 6.5% leukocytes and the CD16^−^ granulocytes comprising a median of 1.3% of leukocytes, however we observed a high level of variability in these cell populations between samples. CD16^+^ granulocytes in BAL have previously been described as neutrophils, whilst CD16^−^ granulocytes have been described as eosinophils. Monocytes, dendritic cells, CD8 T cells, CD4 T cells, B cells and NK cells were also identified in BAL, totalling to a median of 4.6% of leukocytes in BAL combined. The median cell proportions for each of these cell types (expressed as percentage of leukocytes) were 2.14%, 0.63%, 0.77%, 0.88%, 0.15% and 0.10%, respectively (Figure 1A).

For whole blood, neutrophils comprised a median of 25.8% of CD45^+^ leukocytes while eosinophils were 1%. As expected, CD4 T cells, CD8 T cells, B cells and NK cells existed at much higher frequencies in blood relative to BAL, circulating at a median 20.4%, 9.9%, 9.3% and 2.9% of blood leukocytes, respectively. Blood monocytes could be subtyped into classical, intermediate and non-classical subsets based on CD14 and CD16 expression, with classical monocytes the most abundant (2.4% of leukocytes), followed by non-classical (0.4%) and intermediate (0.24%) monocytes. This is distinct from BAL, where only CD16^−^ (classical) and CD16^+^ (intermediate) subsets could be identified (Figure 1A, Supplementary Figure 1). CD11c^+^ myeloid dendritic cells were also identified in whole blood, comprising a median 0.22% of leukocytes (Figure 1B).

### Immune cell profiling of BAL and whole blood by unsupervised analysis

To confirm the findings observed by manual gating and explore the immune profile of our sample types in more detail, we performed unsupervised clustering analysis of BAL and whole blood samples using FlowSOM (Van Gassen et al., 2015). To visualise these data in two dimensions, the non-linear dimensionality reduction technique UMAP (Konopka, 2018) was applied and the cells colour highlighted by their respective FlowSOM cluster (Figure 1C-F).

Clustering analysis of BAL samples with FlowSOM using 13 lineage markers revealed 15 cell clusters (Figure 1C). Based on the expression of lineage markers, the clusters were classified into annotated cell types. This revealed clusters corresponding to four alveolar macrophage populations, four granulocyte populations, an undefined immature macrophage-like population, and one cluster corresponding to each of NK cells, CD8 T cells, CD4 T cells, dendritic cells, CD16^−^ monocytes, and CD16^+^ monocytes. The average frequencies of each cluster across all BAL samples are shown in Figure 1C. Clusters corresponding to alveolar macrophage populations were highly positive for CD206, CD11c and HLA-DR, whilst demonstrating an intrinsically auto-fluorescent signature, as previously reported (Pons et al., 2005). Clusters corresponding to granulocyte populations all expressed CD15 and CD66b, with granulocyte clusters 1 and 2 also expressing HLA-DR, and the most abundant granulocyte cluster (number 4) expressing high levels of CD16. Monocyte clusters expressed characteristic markers such as CD11c, HLA-DR and CD14, however the CD16^+^ monocyte population also expressed CD206. To further verify our clustering results, the UMAP analysis depicted in Figure 1D shows that FlowSOM clusters corresponding to macrophage populations, granulocyte populations, monocyte populations, dendritic cells, NK cells, CD4 T cells and CD8 T cells cluster together as expected within two-dimensional space and correspond well to the automatically defined UMAP clusters.

Clustering analysis of whole blood samples with FlowSOM using the same 13 lineage markers also revealed 15 cell clusters (Figure 1E). Based on the expression of lineage markers, the clusters were classified into annotated cell types. This revealed clusters corresponding to three neutrophil populations, two monocyte populations (CD16^−^ and CD16^+^), two NK cell populations (CD16^−^ and CD16^+^), two B cell populations, an undefined granulocyte-population, and one cluster corresponding to each of eosinophils, CD8 T cells and CD4 T cells. Two clusters (totalling to 2.3%) were unable to be confidently classified. The average frequencies of each cluster across all whole blood samples are shown in Figure 1E. Clusters corresponding to neutrophils expressed high levels of CD15 and CD16, whilst eosinophils expressed low levels of CD15 and were positive for HLA-DR and CD66b. The UMAP analysis depicted in Figure 1F shows that FlowSOM clusters corresponding to circulating neutrophils, eosinophils, monocytes, B cells, NK cells, CD4 T cells and CD8 T cells cluster together as expected within two-dimensional space, corresponding well to the automatically defined UMAP clusters.

For both BAL and whole blood, frequencies of major cell types identified by unsupervised analysis were statistically significantly comparable to those obtained by manual gating for each individual (Supplementary Figures 3-4).

### Relationship between immune cell profiles across lung and blood compartments

As we generated flow cytometry data on matched lung and blood samples from every individual, we next explored the relationship between immune cell profiles within and across the two compartments. As can be seen in Figure 1G, granulocytes from BAL expressed higher levels of CD66b, CD11b and CD15 relative to the equivalent granulocyte population in whole blood. Similarly, CD16^+^ monocytes from BAL expressed high levels of CD206, whilst the CD16^+^ monocytes in blood do not express this marker (Figure 1G).

The heatmap in Figure 1H depicts the immune cell frequencies in BAL and blood from the 11 participants included in this study. We observed a positive correlation between the proportion of BAL neutrophils and eosinophils (r=0.85, p=0.001), BAL CD4 T cells and CD8 T cells (r=0.86, p=0.004), as well as BAL monocytes and dendritic cells (r=0.7, p=0.020). Conversely, a negative correlation between the proportion of BAL alveolar macrophages and neutrophils (r=-0.89, p=0.0005), as well as BAL alveolar macrophages and eosinophils (r=-0.66, p=0.03), was also shown. We additionally observed a positive correlation between the proportion of BAL monocytes and circulating blood dendritic cells (r=0.62, p=0.047) (Figure 1I).

## DISCUSSION

In this work, we developed new experimental methods and applied unsupervised analytical tools to profile the local and systemic immune response in childhood respiratory disease. We comprehensively profiled immune cell populations across blood and lung compartments in children, showed that inflammatory cells in the BAL express higher levels of activation and migration markers, and applied unsupervised analytical tools to reveal novel monocyte and macrophage populations in the paediatric lung. While these data were derived only from children with CF, the methods described could be applied to any childhood respiratory disease and will assist in improving our understanding of immunity and inflammation in these conditions.

The most common immune cell population in the lung is the alveolar macrophage, comprising up to 90% of immune cells in the bronchoalveolar lavage fluid in children, shown by us previously as well as in the current study. Recent work has also showed distinct transcriptional profiles in macrophages obtained from fetal and adult lungs (Bian et al., 2020), highlighting that alveolar macrophages undergo significant development throughout life. Another recent finding revealed that in adults who have undergone lung transplantation, the majority of alveolar macrophages are derived from the recipient’s bone marrow, rather than from self-replenishing resident populations in the donor lung (Byrne et al., 2020). Furthermore, sc-RNAseq analysis of the adult lung identified multiple alveolar macrophage subpopulations (Bassler et al., 2020), a novel finding yet to be explored in children. Our unsupervised hierarchical clustering analysis revealed four alveolar macrophage subpopulations based on differences in expression of lineage markers. Future work will investigate functional differences between these subpopulations.

The value of understanding alveolar macrophages was also recently highlighted by Liao *et al*, who examined the sc-RNAseq profile of adults with COVID-19 infection, demonstrating that the alveolar macrophage subtype composition in BAL fluid was associated with disease severity, and that those with severe disease had higher proportions of migratory monocyte-derived macrophages which secreted IL-6 (Liao et al., 2020). This highlighted IL-6 as a therapeutic target in severe COVID-19 infection, which was supported by preliminary observational studies showing that IL-6 blockade with monoclonal antibodies improved outcomes (McCreary and Meyer, 2021). This finding has since been confirmed in larger multinational trials.(Gordon et al., 2021) In this setting, a betting understanding of the cell composition in the lung identified potential prognostic biomarkers and therapeutic targets worthy of further evaluation.

Along with alveolar macrophages, granulocyte populations play a key role in driving inflammation in the lung. These data were derived from children with CF in whom granulocytic inflammation is a key driver of disease severity. In a seminal study, Sly *et al* demonstrated the presence of free neutrophil elastase in early life BAL was the best predictor of future lung disease severity.(Sly et al., 2013) In the current work, we show that BAL clusters corresponding to granulocyte populations all expressed CD15 and CD66b, with varying expression of HLA-DR and CD16. When compared to blood granulocytes, BAL granulocytes showed higher median levels of CD66b, CD11b and CD15, all markers involved in activation and migration of granulocytes in the lung (Fortunati et al., 2009; Kargl et al., 2017).

We also show that monocytes make up a small proportion of immune cells in the lung and consist of two primary populations (CD16^+^ and CD16^−^). The CD16^−^ monocytes expressed the alveolar macrophage marker CD206, a feature not observed on blood monocytes. We further revealed a monocyte/macrophage-like cluster that demonstrated some of the auto-fluorescent signatures of alveolar macrophages. Collectively, these findings suggest that in addition to the tissue-resident alveolar macrophages, there exists a population of monocyte-derived macrophages within the lung of children with CF. Previous studies have shown that circulating monocytes are recruited to the lung during disease to orchestrate a pro-inflammatory immune response (McQuattie-Pimentel et al., 2018). Whether monocyte-derived macrophages are causally involved in CF inflammation will be area of future investigation. In a study of adults with COPD, peripheral blood monocytes shared an overlapping gene signature with alveolar macrophages. This signature correlated with lung function, highlighting that circulating monocytes and alveolar macrophages may both be involved in disease progression (Poliska et al., 2011).

There are several limitations related to these data that must be acknowledged. Firstly, these data are derived exclusively from children with CF which may limit the applicability to other conditions. In addition, the flow cytometry panel used will influence the number and nature of cell subtypes identified. It is possible that with a broader panel of markers, a larger number of cell subtypes would have been identified. Lastly, these data do not provide insight into the function or clinical impact of the cell subtypes described. Despite these limitations however, the work achieves its primary aim of developing advanced experimental and analytical methods for immune cell profiling of early life respiratory disease. In particular, the methods highlight that it is possible to obtain and utilise samples collected at the time of clinically performed procedures for downstream analysis to better understand inflammation in childhood respiratory diseases. While only children with CF were included in the current study, the same methods could be used to explore other common childhood diseases. In addition, the work highlights the value to augmenting traditional manual gating-based analysis of flow cytometry data with unsupervised analysis. The use of FlowSOM and UMAP in analysing the current data allowed identification of several new cell subtypes.

There are several future directions that will stem from this work. Further experiments will characterise the functional impact of the new cell types identified. In the first instance, fluorescence-activated cell sorting can be used to purify the subtypes and then they can be evaluated using *in vitro* stimulation experiments. In parallel to this, by employing these techniques on a larger number of samples and comparing cell type proportions to clinical outcomes (such as severity of structural lung disease measured by computed tomography), the relationship between cell subtypes and clinical disease severity can be explored. Lastly, simultaneous analysis of blood and BAL with other single cell techniques at a transcriptomic (CITE-seq) or epigenomic level (sc-ATAC-seq) would further improve our understanding of the mechanisms underlying childhood respiratory diseases.

In conclusion, the methods described in this study permit improved immune cell profiling and analysis of childhood respiratory diseases. These techniques can be applied to a number of childhood respiratory diseases with a potential translational impact via the identification of clinical biomarkers and therapeutic targets.

## Acknowledgements

M.R.N is supported by a Melbourne Children’s LifeCourse Fellowship. This work was supported by a Vertex Innovation Award by Vertex Pharmaceuticals. Vertex did not have any input into study design, study execution, analysis of results or manuscript development. We thank the children and parents who participate in the AREST-CF study.

