## Supplemental data for "Application of high dimensional flow cytometry and unsupervised analysis to define the immune cell landscape of early childhood respiratory and blood compartments"

**Supp Table 1.** Flow cytometry antibody cocktail for BAL and blood samples

| **Surface Marker** | **Fluorophore** | **Clone** | **Final Dilution** |
|---|---|---|---|
| CD14 | BV786 | M5E2 | 1:50 |
| CD11b | BUV805 | ICRF44 | 1:100 |
| CD45 | BV711 | HI30 | 1:100 |
| CD56 | BUV737 | NCAM16.2 | 1:100 |
| CD11c | PE-Cy7 | B-ly6 | 1:100 |
| CD63 | A647 | H5C6 | 1:100 |
| CD4 | A700 | RPA-T4 | 1:100 |
| CD3 | BB515 | UCHTI | 1:100 |
| CD15 | PE-CF594 | W6D3 | 1:200 |
| HLADR | V500 | G46-6 | 1:200 |
| CD19 | BV605 | SJ25C1 | 1:200 |
| CD8 | BV650 | RPA-T8 | 1:200 |
| CD206 | BV421 | G10F5 | 1:200 |
| CD66b | PE | 19.2 | 1:200 |
| CD16 | BUV395 | 3G8 | 1:400 |
| Live/dead | N-IR | | |

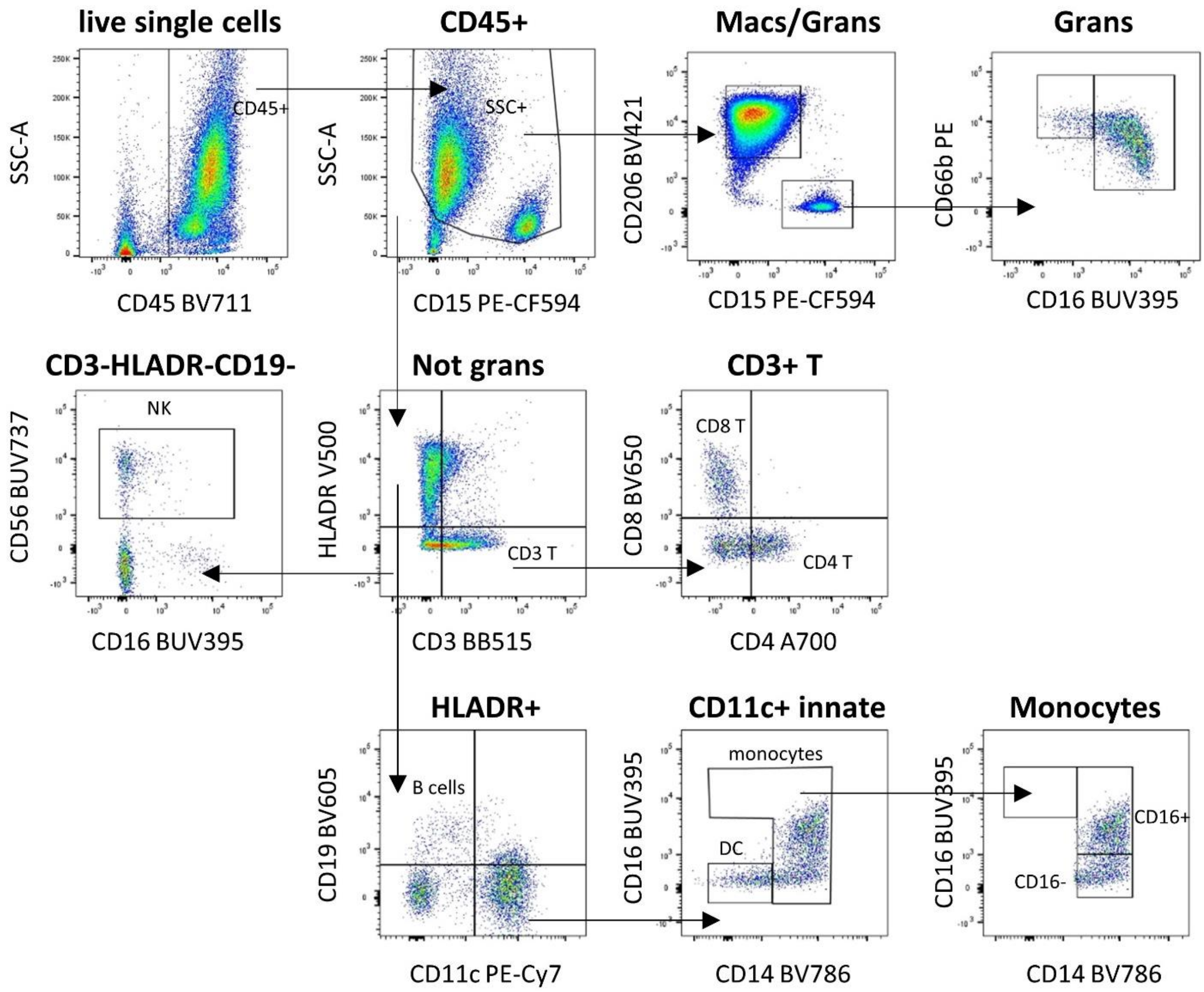


**Supp Figure 1. Representative flow cytometry gating strategies for BAL.** CD45+ leukocytes were firstly selected from the live single cell fraction. Within CD45+ cells, macrophages/granulocytes were selected based on a high SSC-A profile. Macrophages were identified based on $CD206^{+}CD15^{-}$ phenotype, whilst granulocytes were $CD206^{-}CD15^{+}$. Within the granulocyte fraction, $CD16^{+}$ and $CD16^{-}$ granulocytes were identified. $CD45^{+}SSC\text{-}A^{low}$ cells were further subtyped into $HLADR^{+}$ and $CD3^{+}$ T cells. Within the $CD3^{+}$ T cell fraction, CD4 and CD8 T cells were identified. $HLADR^{+}CD19^{+}$ cells were identified as B cells, and $HLADR^{+}CD11c^{+}$ cells were identified as innate cells. Within the innate cell fraction, monocytes were selected based on CD14 expression, whilst DCs were $HLADR^{+}CD11c^{+}CD14^{-}$ $CD16^{-}$. Monocytes were also assessed for CD16 expression, revealing both $CD16^{+}$ and $CD16^{-}$ subsets. $CD3^{-}HLADR^{-}CD19^{-}CD56^{+}$ cells were identified as NK cells.

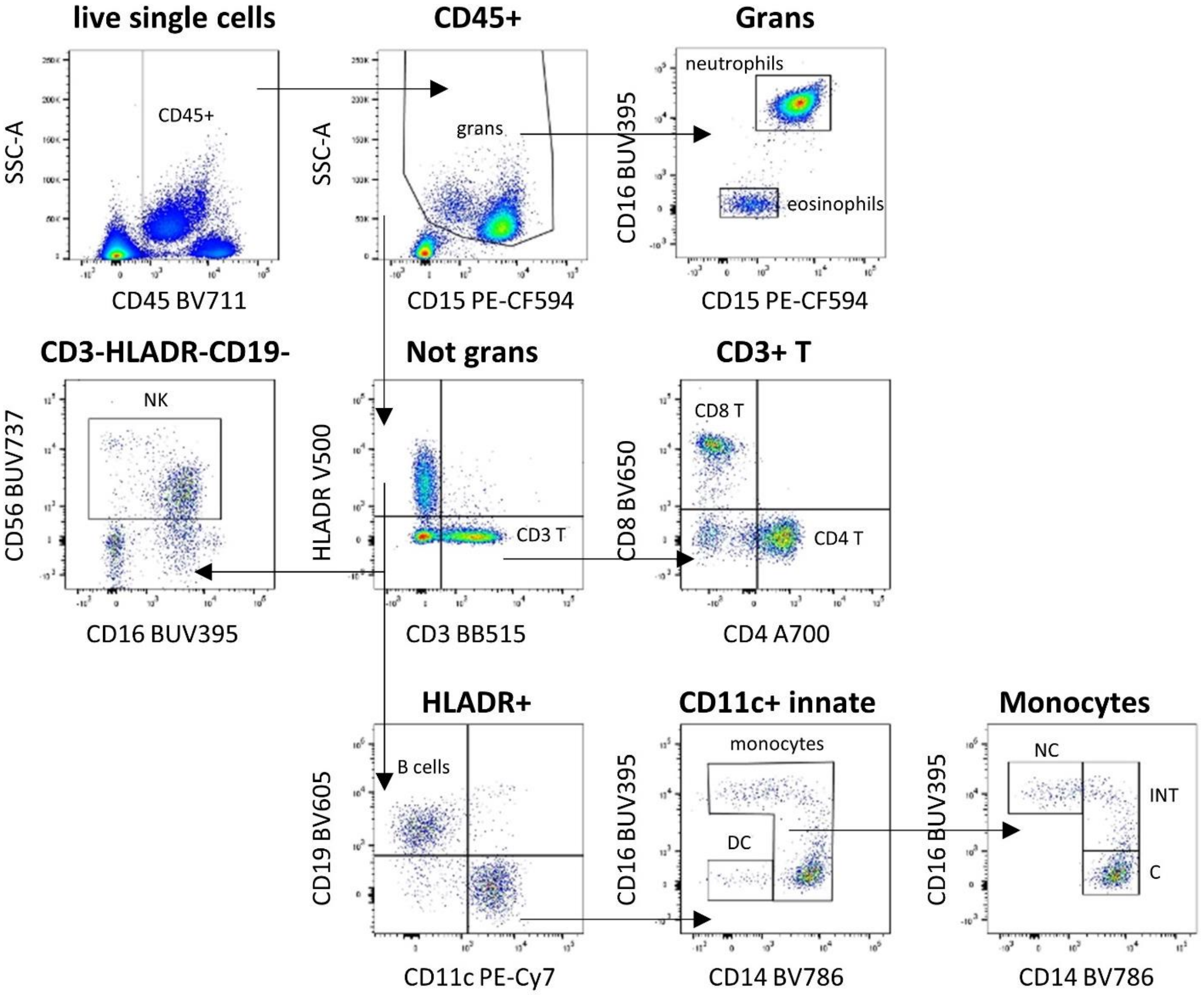


**Supp Figure 2. Representative flow cytometry gating strategies for whole blood.** CD45+ leukocytes were firstly selected from the live single cell fraction. Within CD45+ cells, granulocytes were selected based on $CD15^{+}$ and $SSC\text{-}A\text{-}^{high}$ profile. Within the granulocyte fraction, neutrophils were identified based on $CD15^{high}CD16^{+}$ phenotype, whilst eosinophils were $CD15^{low}CD16^{-}$. $CD45^{+}SSC\text{-}A^{low}$ cells were further subtyped into $HLADR^{+}$ and $CD3^{+}$ T cells. Within the $CD3^{+}$ T cell fraction, CD4 and CD8 T cells were identified. $HLADR^{+}CD19^{+}$ cells were identified as B cells, and $HLADR^{+}CD11c^{+}$ cells were identified as innate cells. Within the innate cell fraction, monocytes were selected based on CD14 expression, whilst DCs were $HLADR^{+}CD11c^{+}CD14^{-}CD16^{-}$. Monocytes were also assessed for CD16 expression, revealing three subsets: non-classical ($CD14^{low}CD16^{+}$), intermediate ($CD14^{+}CD16^{+}$) and classical ($CD14^{+}CD16^{-}$). $CD3^{-}HLADR^{-}CD19^{-}CD56^{+}$ cells were identified as NK cells.

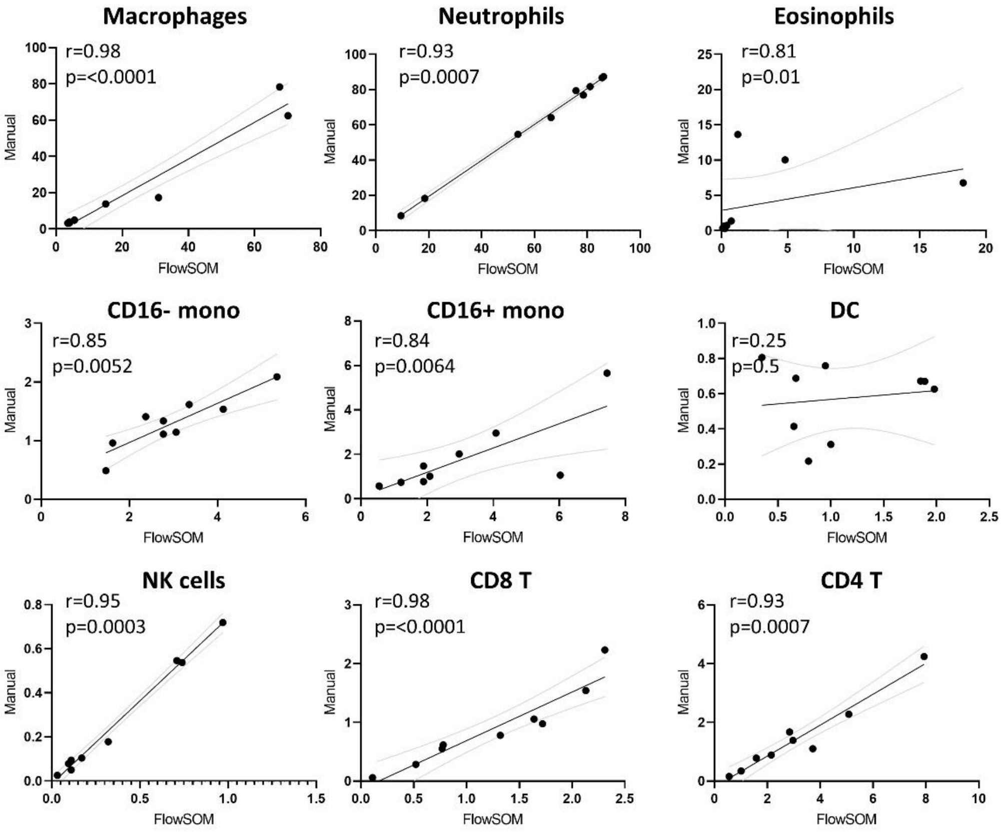


**Supplementary Figure 3.** Correlation of cell type proportions in BAL obtained from unsupervised clustering (FlowSOM) and manual gating. Percentages represent the % live $CD45^+$ cells. Correlation co-efficient and p values are from two-tailed spearman tests.

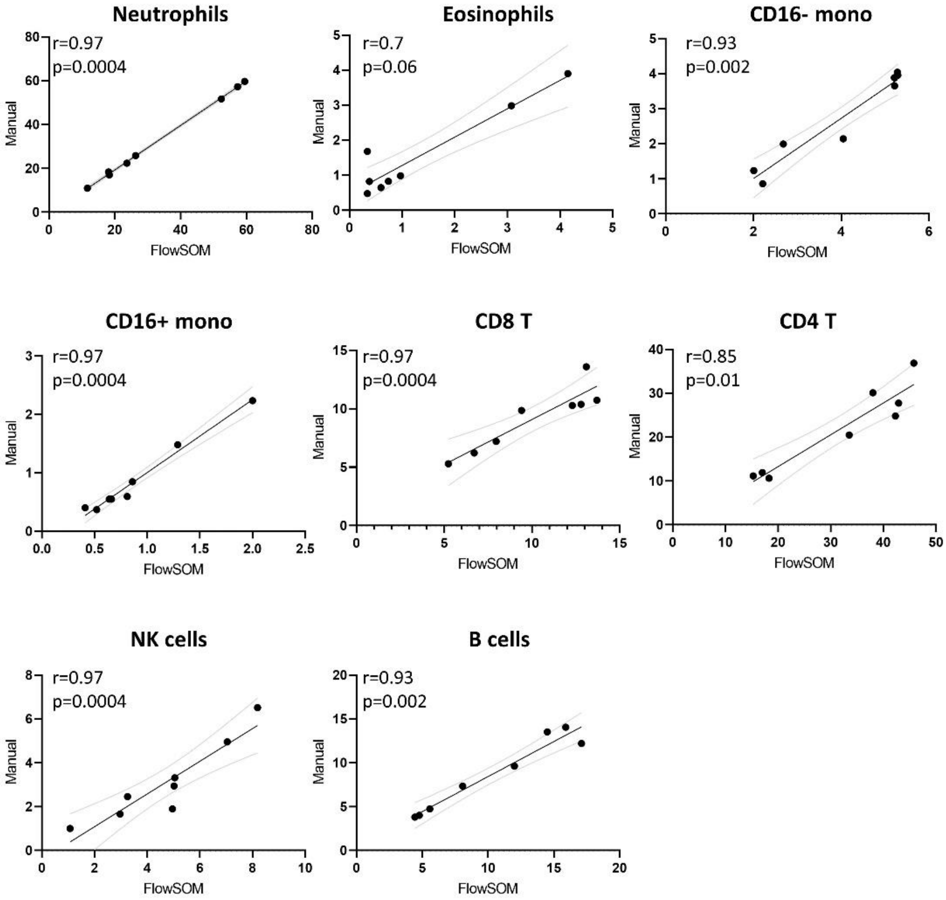


**Supplementary Figure 4.** Correlation of cell type proportions in blood obtained from unsupervised clustering (FlowSOM) and manual gating. Percentages represent the % live $CD45^+$ cells. Correlation co-efficient and p values are from two-tailed spearman tests.
